# Functional Genetic Screen to Identify Interneurons Governing Behaviorally Distinct Aspects of *Drosophila* Larval Motor Programs

**DOI:** 10.1101/041061

**Authors:** Matt Q. Clark, Stephanie J. McCumsey, Sereno Lopez-Darwin, Ellie S. Heckscher, Chris Q. Doe

## Abstract

Drosophila larval crawling is an attractive system to study patterned motor output at the level of animal behavior. Larval crawling consists of waves of muscle contractions generating forward or reverse locomotion. In addition, larvae undergo additional behaviors including head casts, turning, and feeding. It is likely that some neurons are used in all these behaviors (e.g. motor neurons), but the identity (or even existence) of neurons dedicated to specific aspects of behavior is unclear. To identify neurons that regulate specific aspects of larval locomotion, we performed a genetic screen to identify neurons that, when activated, could elicit distinct motor programs. We used 165 Janelia *CRM-Gal4* lines – chosen for sparse neuronal expression – to express the warmth-inducible neuronal activator TrpA1 and screened for locomotor defects. The primary screen measured forward locomotion velocity, and we identified 63 lines that had locomotion velocities significantly slower than controls following TrpA1 activation (28°C). A secondary screen was performed on these lines, revealing multiple discrete behavioral phenotypes including slow forward locomotion, excessive reverse locomotion, excessive turning, excessive feeding, immobile, rigid paralysis, and delayed paralysis. While many of the Gal4 lines had motor, sensory, or muscle expression that may account for some or all of the phenotype, some lines showed specific expression in a sparse pattern of interneurons. Our results show that distinct motor programs utilize distinct subsets of interneurons, and provide an entry point for characterizing interneurons governing different elements of the larval motor program.

## Introduction

Understanding the neurobiological basis of behavior and brain disorders is a grand challenge of the 21^st^ century as outlined by the BRAIN Initiative [1]. The study of invertebrates has yielded numerous insights into the neural basis of behavior [2]. Invertebrates offer an elegant platform to investigate behavioral patterns due to the stereotypy of behaviors as well as the ability to reproducibly identify individual neurons that generate behaviors. Examples include detailed studies of escape behaviors driven by command neurons of crayfish [3], central pattern generating circuits of crustaceans [4], reciprocal inhibition motifs in the visual system of the horseshoe crabs [5,6], and learning and memory habituation in the sea hare [7]. While these principles were discovered in invertebrates, they are broadly applicable to aspects of neural circuit function in vertebrates.

An integral component of all of motor systems are central pattern generators (CPGs), which underlie the generation of rhythmic motor patterns [8,9]. CPGs are diverse and modular and can be recruited to function depending on context and exposure to aminergic neuromodulators such as serotonin [10]. Neural circuits that comprise CPGs can function autonomous of sensory or descending inputs [11]. The study of insects has led to advances in understanding unique aspects of motor programs including patterned motor output, sensory or descending inputs and the local control of musculature [12,13].

Although it is possible to study neural circuits in *Drosophila melanogaster* [14–17], historically it has been challenging due to the small size and inaccessibility of their neurons. However, the recent advent of advanced techniques to target, label and monitor physiological input and output has made *Drosophila* an excellent model to investigate the neurobiological basis of behaviors and the development of neural circuits [18–24]. Furthermore, serial section Transmission Electron Microscopy (ssTEM) maps of neural connectivity [25–31] and advanced computational ‘ethomic’ approaches to establish behavioral categories [32–34] will greatly aid future investigations.

With approximately 10,000-15,000 neurons [35], *Drosophila* larvae offer a relatively simple preparation for investigating neural circuit formation at single cell resolution. Considerable progress has been made in understanding larval and embryonic neurogenesis with markers of neuroblasts and progeny well characterized [36–39]. Recent anatomical studies show that many, if not all, interneurons of the ventral nerve cord (VNC) have a unique morphology [40], and possibly unique molecular profile [41]. Importantly, there are over 7,000 Gal4 lines generated by the Rubin lab [42]; we previously screened these lines for late embryonic expression, and identified several hundred expressed in sparse numbers of neurons within the VNC [43]. These tools allow us and other researchers genetic access to the majority of interneurons within the VNC, and allow us to characterize their role in late embryonic or newly hatched larval behaviors by expression of ion channels to silence neuronal activity (KiR; [44]) or induce neuronal activity (TrpA1; [24]). By screening these Gal4 patterns for unique behavioral phenotypes it becomes possible to connect neuronal anatomy to neuronal function and development. Recent work in adults has used this approach to connect adult behaviors to their neurogenic origins in the late larva [38].

*Drosophila* larval locomotion is an excellent model to study patterned rhythmicity. Stereotypic movements include turns, head sweeps, pauses, and forward and backwards locomotion (Figure 1A) [45]. Larval forward and reverse locomotion is generated by abdominal somatic body wall muscle contractions moving from posterior to anterior (forward locomotion) or anterior to posterior (reverse locomotion) [46]. Consecutive bouts of forwards or backwards waves are called runs (Figure 1B). Asymmetric contractions of thoracic body wall musculature generate turns [47]. Neural control of turning movements is located within the thoracic segments of the VNC [48] while the CPGs that drive larval locomotion have also been shown to be located in the thoracic and abdominal segments of the VNC [49,50]. However, the specific neurons that comprise the CPG are currently unknown [51]. Similarly, little is known about the neurons specifically used in other aspects of locomotion, such as forward or reverse movements, head sweeps, and pauses.

**Figure 1.**
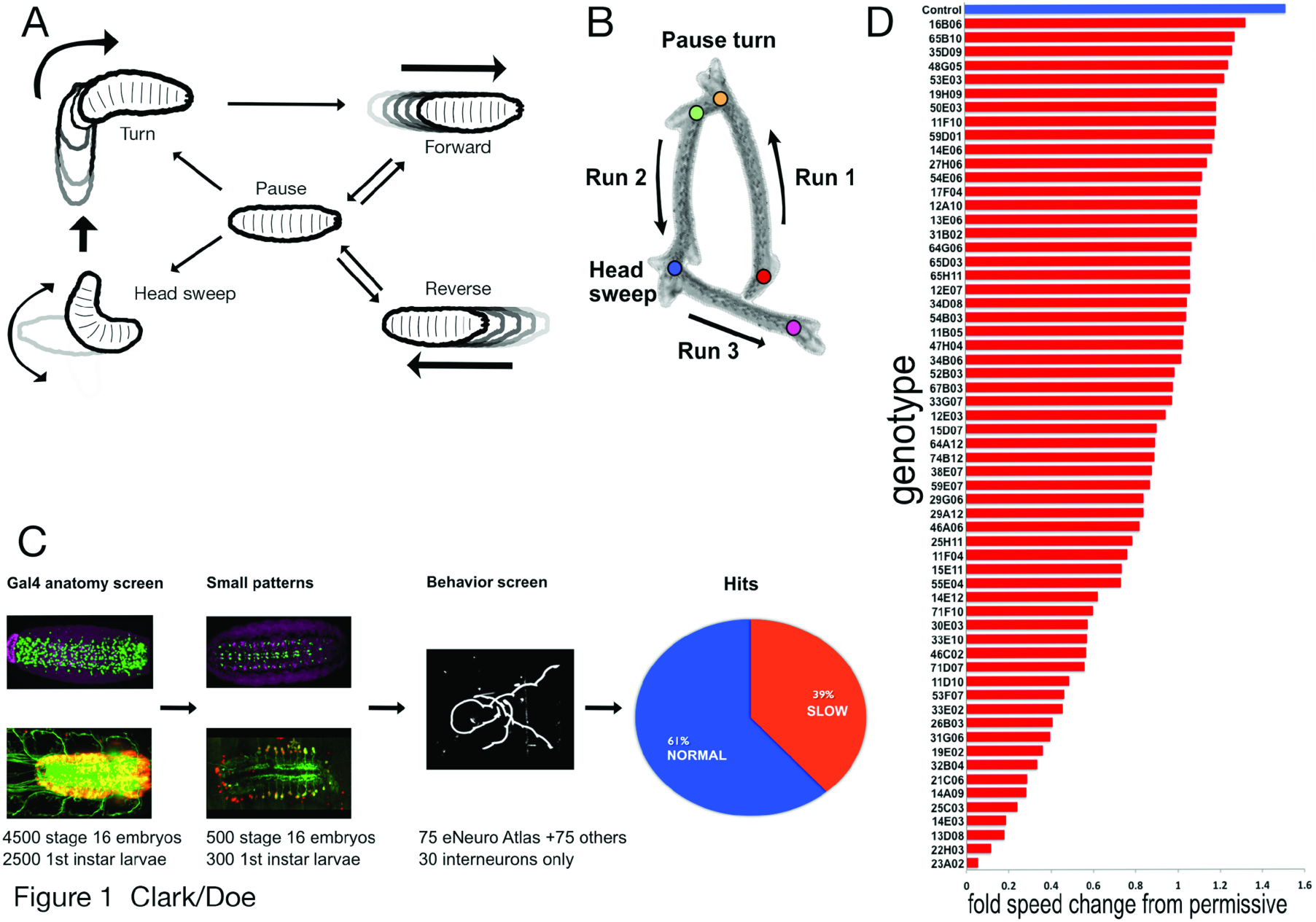
TrpA1 functional screen results and low magnification traces of crawl patterns. **(A)** Ethogram of common behaviors during crawling (modified from reference 52). **(B)** A time-lapse projection of a typical larval crawl pattern consisting of runs, pause turns and head sweeps. **(C)** Initial screening of over 7000 Gal4 patterns yielded at least 700 Gal4 patterns with <15 neurons per hemisegment. 75 of these late stage embryonic Gal4 patterns were entered into eNeuro atlas and screened at first larval instar with ectopically expressed warm-gated cation channel UAS-TrpA1. An additional 100 *CRM-Gal4* expression patterns were screened with TrpA1 resulting in nearly 40% of those exhibiting crawl defects as shown in histogram of speed tracking. **(D)** Tracking speed changes from permissive (23°C) to restrictive (28°C) yielded genotype-specific fold changes statistically slower when compared to controls (top blue). p-values for all represented in red was <0.05 (student’s t-test).

Here we screen a collection of several hundred Gal4 lines that are sparsely expressed in the CNS to identify neurons that, when activated, can induce specific alterations in the larval locomotor program. The results presented here will provide the basis for future functional studies of motor control and neural circuit formation in the *Drosophila* larva.

## Materials and Methods

### Imaging Gal4 expression patterns in whole first instar larvae

For every Gal4 line we imaged whole newly-hatched “L0” first instar larvae, defined as between 0-4 hours of hatching, for native GFP fluorescence and nuclear red stinger fluorescence. We used a newly developed protocol to fix and stain intact larvae to confirm the expression pattern. Briefly, intact L0-L3 larvae were prepared for staining by incubation in 100% bleach for 10 min at room temperature (rt), digested with chymotrypsin/collagenase for 1 h at 37C, fixed in 9% formaldehyde for 30 min at rt, incubated in 1:1 methanol:heptane for 1 min at rt, and post-fixed in methanol for 1-3 days at −20C (L. Manning and CQD, in preparation). Subsequently, standard methods were used for staining with chick anti-GFP (1:2000; Aves) [43].

### Bright-field whole larva behavioral recordings

All behavior was done using “L0” first instar larvae. Behavior arenas were made of 6% agar in grape juice, 2 mm thick and in 5.5 cm in diameter. Temperature was measured using an Omega HH508 thermometer with type K hypodermic thermocouple directly measuring agar surface temperature. Temperature was controlled using a custom-built thermoelectric controller and peltier device. The arenas were placed under a Leica S8APO dissecting microscope and red light (700 nm, Metaphase Technologies) illuminated a single larva. The microscope was equipped with a Scion1394 monochrome CCD Camera, using Scion VisiCapture software. Images were acquired via ImageJ at either 4 Hz for low magnification videos or 7.5 Hz for high magnification.

### TrpA1 screen

Adult UAS-TrpA1 virgin females were crossed to males of select Janelia *CRM-Gal4* lines which were kept in standard collection bottles (Genesee Scientific) and allowed to lay eggs on apple caps with yeast paste. For low magnification screening, a single larva was staged on a behavior arena and given a 5-10 minute period of acclimation. For recordings, larvae were permitted to crawl freely, and the stage was manually re-centered when the larva left the field of view. Individual larvae were recorded at permissive (23°C) and restrictive (28°C) for 800 frames at 4 Hz.

### Quantification of crawl parameters

We conducted two locomotion assays, low magnification for screening and high magnification in order to discern the etiology of crawl defects. For our initial low magnification screening, we calculated the speed of larval locomotion with automated analysis using custom Matlab scripts. Scripts were written in MATLAB and are available upon request.

#### Object recognition

For low magnification tracking an individual larva was detected in each frame using the following steps. The image was mildly blurred using a Gaussian blurring function to reduce background artifacts and make the appearance of the larva more uniform. The built-in MATLAB thresholding function utilizing Otsu’s method was used to segment the image. The image was then made binary and objects were morphologically closed. In each frame a single object was selected as the larva based on empirically determined and manually entered size. Built-in MATLAB functions were used to determine the larval object’s area and centroid position in each frame. The script returned no data if more than one object was found or no object was found.

#### Crawling Speed

An approximate instantaneous speed was calculated by taking the distance traveled by the larval object between two consecutive frames and dividing by the time elapsed. All instantaneous speeds were then averaged to get an average crawling speed. If there was more than one behavioral recording for a given larva, data from up to three recordings were included. Standard deviation was then calculated. To exclude time points in which the larva appeared to travel large distances due to manual repositioning of larva during behavioral recording, if the distance traveled by the larval object between successive frames was farther than half the length of the larva (see below) then the frames were excluded from speed calculations.

#### Larval length

The mean area of the larva was averaged to get “LarvalLen”; then larval length was calculated as = sqrt(LarvalLen/3.14).

#### Normalized data

Normalized values (*n*) refer to values for a given larva at restrictive (*r*) temperature less the values for that larva at permissive *(p)* temperature, divided by values at permissive temperature *(n = (r-p)/p).*

#### Test statistics

Built-in MATLAB function was used to run a 1-tailed, t-test assuming equal means but unequal variance (‘ttest2’ function).

#### Representation of slow hits

To represent lines that exhibited crawling defects at restrictive temperature, we chose two criteria to define slow crawls. First were those that were slow at restrictive compared to controls (students t-test) and second those that did not increase their speed by the same rate when shifted from permissive to restrictive when compared to control (students t-test). Then average speed at restrictive was then divided by permissive.

#### High mag quantification

We manually calculated head sweeps, and forward and reverse wave propagation.

### Fly Stocks

The following stocks obtained from the Bloomington Drosophila Stock Center (NIH P40OD018537) were used in this study: *10xUAS-IVS-myr∷GFP* (BL #32198), *UAS-RedStinger* (BL# 8546), *UAS-TrpA1* (BL #26263), D42-Gal4 (BL #8816), *OK6-Gal4* (Aberle, 2002), *Mef2-Gal4* (BL #27390), *repo-Gal4* (BL #7415), *elav-Gal4* (BL #8760), EL-Gal4 (Fujioka et al., 2003), *RN2-Gal4* (BL #7470), *CQ-Gal4* (BL #7466), *OK371-Gal4* (BL #26160), *GAD1-Gal4* (BL # 51630), *ple-Gal4* (BL# 8848), *trh-Gal4* (BL# 38389), *painless-gal4* (BL# 27894), *iav-Gal4* BL# (52273), *nan-Gal4* (BL #24903), *en-Gal4* (BL #1973), and *pBDP-Gal4.1Uw* in attP2 (gift from B.D. Pfeiffer). Flies were raised on conventional cornmeal agar medium at 25°C.

## Results

### TrpA1 activation of sparse neuronal subsets results in slower, but not faster, larval locomotion

To identify neurons that can generate specific aspects of locomotor behaviors (pause, turn, forward, reverse, etc.), we screened Janelia *CRM-Gal4* lines containing sparse expression patterns at either embryonic stage 16 or in newly hatched “L0” first instar larvae (0-4h after hatching) (Figure 1C). We began with 7000 *CRM-Gal4* patterns; 4500 were screened at embryonic stage 16 with *UAS-nls∷GFP* marking the cell nucleus, and 2500 were screened at first instar with *UAS-myr∷GFP, UAS-redstinger* labeling the cell membrane and cell nucleus. From the initial 4500 we selected 75 patterns that had sparse expression patterns and entered them into the eNeuro atlas [41], which allows us to determine if they are motor neurons, interneurons, or glia. In addition to these 75 lines, we identified an additional 65 lines that had sparse embryonic VNC expression. A final 30 lines with sparse larval (L0) VNC expression were selected from the 2500 first instar expression patterns. We assayed newly hatched L0 larva behavior because it was closest in time to the stage where our Gal4 expression patterns were documented, making it less likely for the pattern to have changed; most embryonic Gal4 patterns are completely different by third larval instar [42,43].

To assess the function of the neurons labeled by each of these Gal4 lines, we screened nearly 200 strains using the warmth-gated neural activator TrpA1 [24]. In our assay regime we monitored crawl speeds of individual newly hatched larvae at permissive temperature (23°C) and then at restrictive temperature (28°C). As with previous behavior experiments using JRC *CRM-Gal4* constructs [34] we used larvae containing the ‘empty’ vector pBDP-Gal4U crossed to UAS-TrpA1 flies as our control; this transgene does not express TrpA1 in the VNC, and larva have normal locomotor velocities (Figure 1D, top). This is an appropriate control as the experimental Gal4 lines from the Rubin collection have a similar genetic background. We noted that control larvae increased their speed from 65.0 μm/sec at permissive temperature (+/- 47.0 SD, n=10) to 98.7 μm/sec at restrictive temperature (+/- 66.3 SD, n=10), or an increase of roughly 1.5 fold (Figure 1D, top).

Approximately 40 percent of lines we screened exhibited elements of crawl defects. We defined a genotype as slow by the following criteria: at restrictive temperature they were slower compared to controls (student t-test p<0.05) and normalized permissive to restrictive change was statistically different (one-tailed student t-test p<0.05). Of those lines that were slow, approximately half had uniquely evocable behaviors that we describe below. We expected to elicit ‘fast’ crawl phenotypes, however, we only detected normal or slow phenotypes.

### TrpA1 activation of sparse neuronal subsets generates multiple, distinct locomotor phenotypes

Control larvae on naturalistic terrain exhibit pauses, head casts, turns and forward and backward locomotion (Figure 1A,B) [45,52], but in our assay they showed a strong bias towards forward locomotion, perhaps due to the temperature shift from 23°C to 28°C [53] (Figure 2A,A’). Each of the *CRM-Gal4 UAS-TrpA1* lines we characterize below has a defect in the frequency or velocity of forward locomotion (Figure 1D, above), and in this section we describe each of the multiple, distinct locomotor phenotypes observed. We present the phenotype of one representative line in Figure 2, larval expression patterns for representative lines in each category are shown in Figure 3, and the cell type expression patterns for all lines in each category are shown in Figure 4.

**Figure 2.**
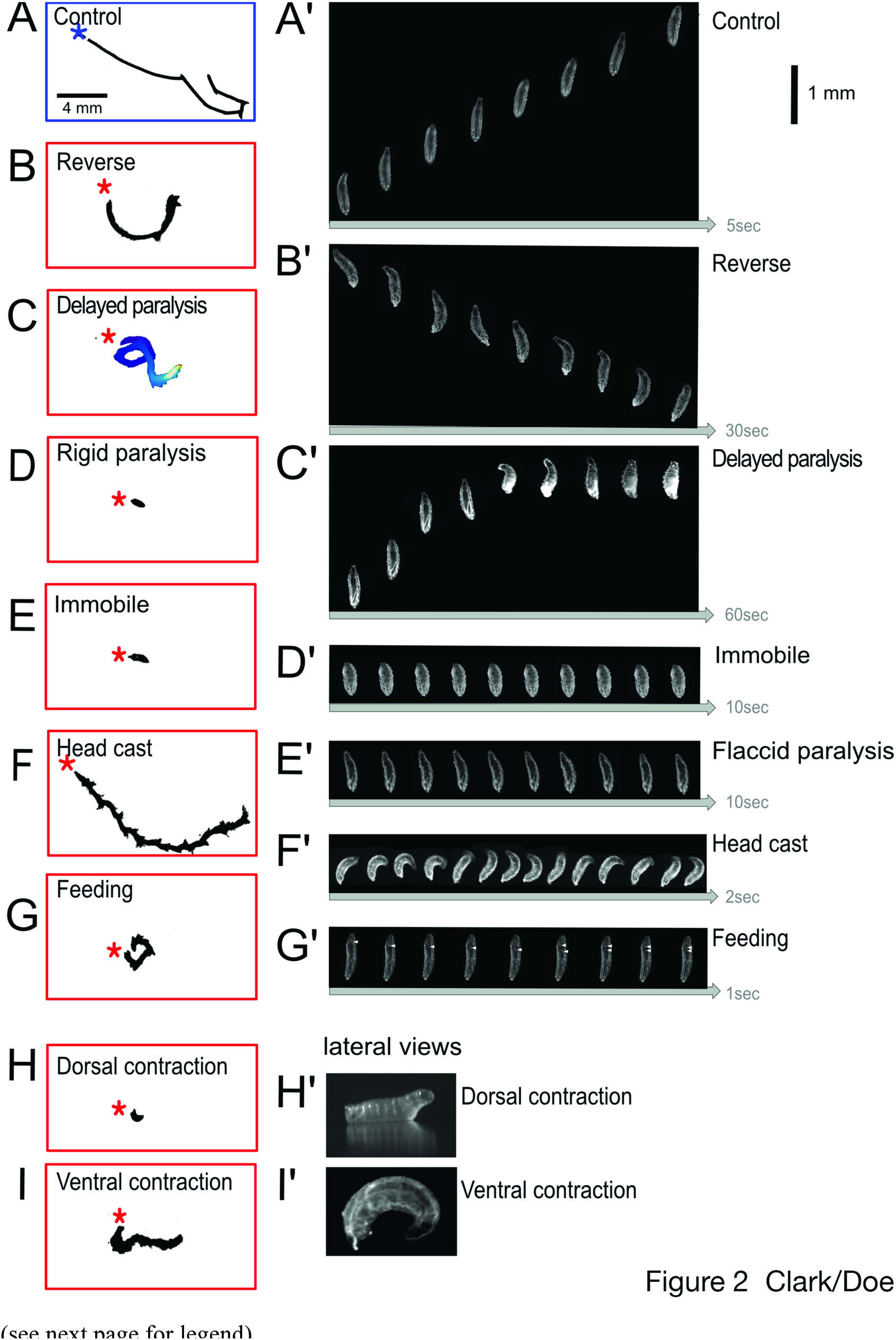
Low and High Magnification Analysis of TrpA1-Induced Crawling Phenotypes. Representational traces of crawl trajectories for control (empty transgene cassette) and TrpA1-induced phenotypes of newly hatched larvae observed at low magnification (left) and high magnification still frames (right). Asterisk denotes beginning of crawl. Still frames from videos of larvae at restrictive temperature were taken at 7.5fps. Phenotype categories are indicated; distance scale bar applies to all right column panels, but each set of movie stills has a unique timeline (arrow at bottom of panel). **(A-A’) Control.** Larva demonstrates a typical crawl with runs and pause turns (left), while larva shown (right) 4 ~travels μM in 5 seconds. **(B-B’) Reverse.** Larva successfully generates complete waves from anterior to posterior only. Translational movements occur strictly in the reverse direction. **(C-C) Delayed paralysis.** Characterized by a free range of movements at restrictive, yet progressively slows until all segments are tonically contracted at 60 seconds. **(D-D’) Rigid paralysis.** All segments are fully contracted with no translational movement. **(E-E’) Immobile.** All segments are fully relaxed with no translational movement. **(F-F’) Head cast.** Crawl trajectory illustrates the ‘back-and-forth’ nature of movement. Peristalsis functions similar to controls, however before a peristaltic wave fully traverses from posterior to anterior, the larva has already begun a head sweep. **(G-G’) Feeding.** Characteristics of ingestion including pharyngeal pumping, mouth hook movement and head tilting. White arrow indicates rhythmic bubble ingestion (larva viewed ventrally). **(H-H’) Dorsal contraction.** Head and tail off the substrate illustrated in lateral view. **(I-I’) Ventral contraction.** Ventral contraction displays little movement and most extreme pictured is stuck ventrally curved. Genotypes: (A) UAS-TrpA1/+; pBDP-Gal4U/+. (B) UAS-TrpA1/+; R53F07-Gal4. (C) UAS-TrpA1/+; R55B12-Gal4/+. (D) UAS-TrpA1/+; R23A02-Gal4. (E) UAS-TrpA1/+; R31G06-Gal4/+. (F) UAS-TrpA1/+; R15D07-Gal4/+. (G) UAS-TrpA1/+; R76F05-Gal4/+. (H) UAS-TrpA1/+; R26B03-Gal4/+. (I) UAS-TrpA1/+; R79E03-Gal4/+.

**Figure 3.**
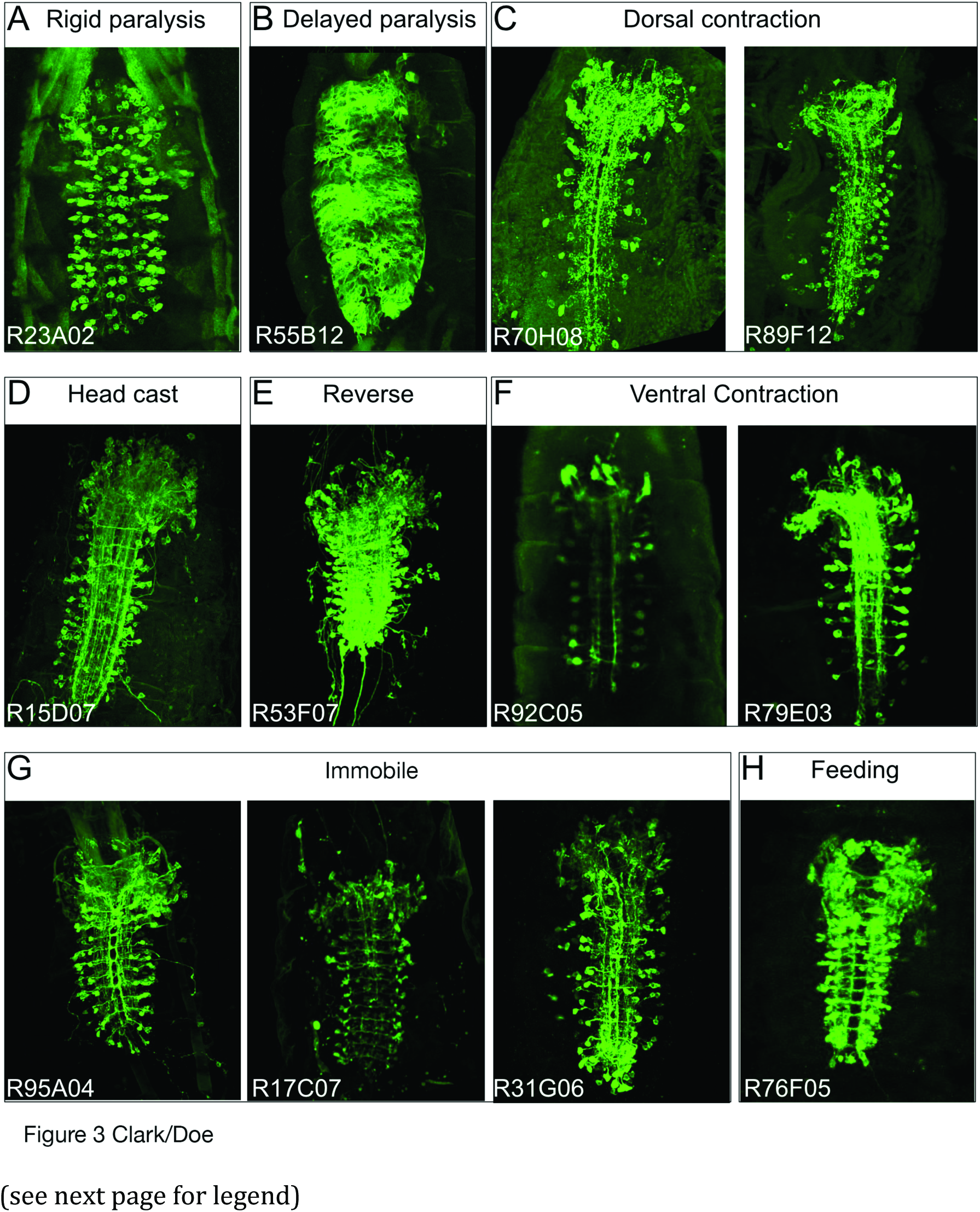
Expression patterns for each phenotype group. Ventral view of Z-stack projections for Gal4 patterns expressing membrane marker UAS-myr∷GFP, except delayed paralysis which is also expressing UAS-redstinger to show cell nucleus. Anterior is up. **(A) Rigid paralysis.** All lines expressed in interneurons and other tissues, with many expressing in all muscles. **(B) Delayed paralysis.** Shown is one slice of z-stack to illustrate the reticulated nature of astrocyte glia in the VNC. Cell nuclei in red. **(C) Dorsal contraction.** Lines shown are interneuron-specific. **(D) Head cast.** This line expresses in interneurons and sporadically in dorsally projecting motor neurons **(E) Reverse.** This line expresses in interneurons and in dorsally projecting motor neurons. **(F) Ventral contraction.** Lines shown are interneuron-specific. **(G) Immobile.** Lines shown are interneuron-specific, with the expression of 31G06 expressed in VO muscles. **(H) Feeding.** One line is interneuron-specific; others express in interneurons as well as motor and sensory neurons.

**Figure 4.**
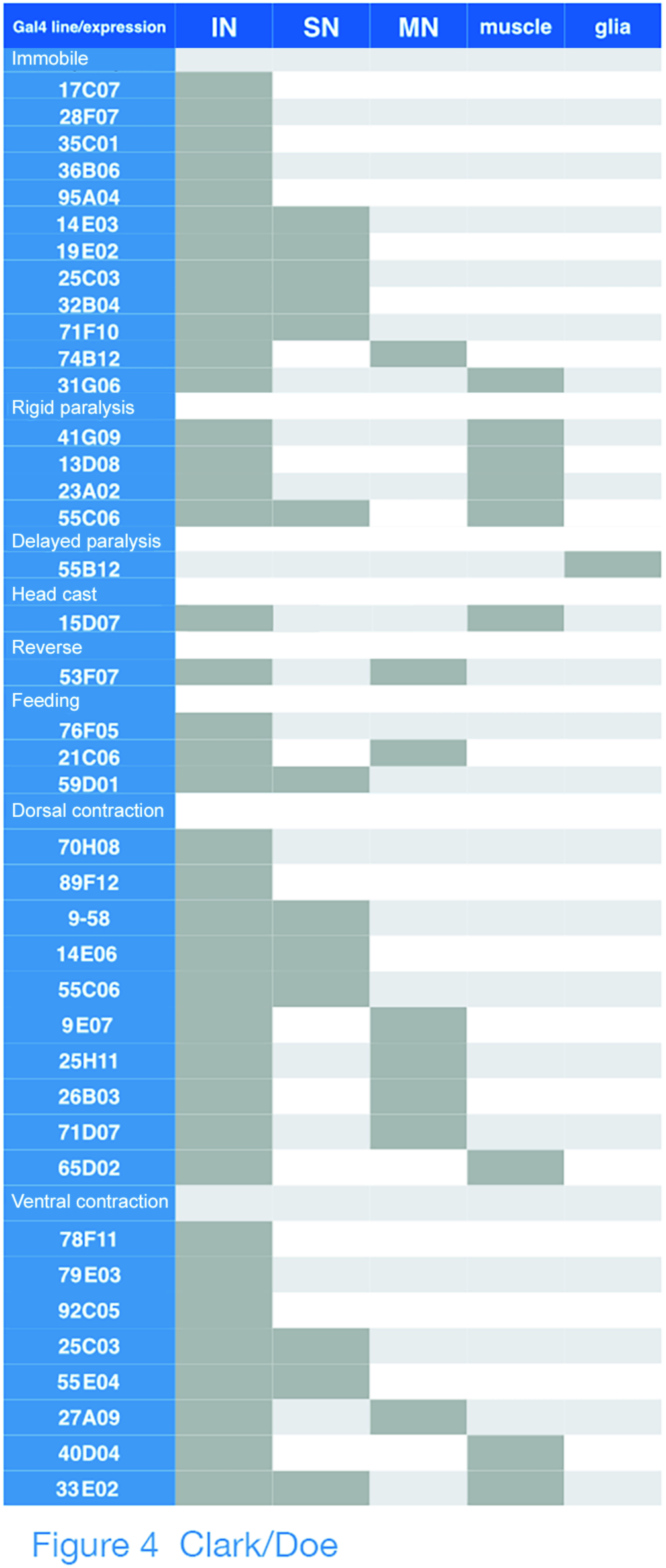
Gal4 line expression patterns in newly hatched larvae. Left column indicates the Janelia Gal4 line name (nomenclature: Rxxxxx) and relevant phenotypic categories. Dark grey boxes to the right indicate cell type expression patterns of each Gal4 line: interneurons (IN), sensory neurons (SN), motor neurons (MN), muscle, and glia.

#### Reverse

We found one line in this category: R53F07 (Figure 2B,B’). Whereas control larvae normally display a range of movements (Figure 1A,B), larvae in this category are strongly biased towards reverse locomotion. Forward propagating waves were generated occasionally, but they often failed to reach the anterior thoracic head region, instead switching prematurely to reverse waves.

Anatomical characterization shows both interneurons and motor neurons (Figure 3E, 4), but many other lines contained motor neurons without showing the reverse locomotion phenotype. We also did not observe expression in any sensory neurons such as the Bolwig organ or Class IV MD neurons, which have been shown to play a role in light mediated aversive response [54]. This suggests that the phenotype is due to activation of one or more interneurons in the pattern.

#### Immobile

We found 12 lines in this category, including R17C07 and 95A04 that showed expression only in interneurons (Figure 2E,E’, 3G). Behavioral hallmarks of this category were loss of mobility with infrequent peristaltic waves. At times, some body wall segments appeared to lack tone and showing a smooth, elongated body shape (Figure 2E’). Larvae could move when prodded, however, distinguishing this category from the next two “paralysis” categories.

Anatomical characterization showed sparse interneuron expression as well as a few lines with additional sensory neuron, motor neuron, or muscle expression (Figure 3G, 4).

#### Rigid paralysis

We found 4 lines in this category, including R23A02 (Figure 2D,D’). Hallmarks of this category include immobility, tonic contraction of all body segments and shortening of larval body length. There was also a nearly complete lack of forward and reverse peristaltic waves. Larvae did not move when prodded.

Anatomical characterization shows lines that contained all body-wall muscles, all motor neurons or large subsets of interneurons (Figure 3A, 4). This last group includes lines that were picked for our behavioral assay due to sparse numbers of interneurons in the late embryo, but ultimately showed greatly increased numbers of interneurons in newly hatched larvae.

#### Delayed paralysis

We found one line in this category: R55B12 (Figure 2E,E’). Larvae appeared identical to controls upon shifting to 28°C, but over time exhibited full tonic contraction paralysis (Figure 2C’). Larvae are sometimes observed recovering from this paralysis but continue to cycle through paralysis periodically. Paralyzed larvae did not move when prodded.

Anatomical characterization showed expression of R55B12 restricted to neuropil “astrocyte” glia. A similar phenotype of “delayed paralysis” was obtained by crossing the glial-specific Repo-Gal4 line to U AS-TrpA1 and shifting to 28°C (data not shown), confirming that the phenotype is due to glial activation.

#### Head cast

We found one line in this category: R15D07 (Figure 2F,F’). Larvae had a “zigzag” pattern of locomotion (Figure 2F) due to persistent head casting (Figure 2F’). Whereas control larvae normally exhibit head casts as part of their exploratory program [55], larvae in this category exhibited continuous head casts during crawls. High magnification time-lapse analysis reveals that posterior to anterior body wall muscle waves characteristic of forward locomotion still occurred in larvae of this category, but the larva often initiated a head cast prior to completion of the wave of muscle contraction (data not shown).

Anatomical characterization showed expression in interneurons in the brain and VNC, plus dorsally projecting motor neurons (Figure 3D, 4). Because other lines contained dorsally projecting motor neurons without showing the head cast phenotype, we suggest the phenotype is due to activation of brain or VNC interneurons.

#### Feeding

We found three lines in this category; line 76F05 is shown in Figure 2G. Hallmarks of this category were a bias towards feeding behavior, including pharyngeal pumping, rhythmic ingestion that can be observed as air bubbles entering the midgut through the esophagus (white triangles, Figure 2G’), and frequent mouth hook movements and head tilting [56,57]. Larvae of one genotype (R21C06) do not move when at restrictive temperature and exhibited elements of the rigid paralysis phenotype while another (R59D01) exhibited a free range a movement while attempting to feed. The genotype expressing only interneurons (76F05) did not move, but showed normal range of motion of the head.

Anatomical characterization showed that all lines had a sparse pattern of interneurons in the brain and VNC (Figure 3H, 4); R21C06 showed additional expression in motor neurons, which is likely to be the cause of the additional rigid paralysis phenotype.

#### Dorsal contraction

We found 10 lines in this category; the R70H08 and R89F12 lines expressing only in sparse interneuronal patterns are shown in Figure 2H. This phenotype is characterized by the most anterior and posterior segments of the larva lifted vertically off the substrate when viewed laterally (Figure 2H’). The phenotype varies in severity with some larvae permanently stuck with their thoracic head region and tail lifted up. At times some continue crawling but periodically become stuck in this position. This phenotype may arise from premotor interneurons stimulating dorsal projecting motor neurons, and we have confirmed that TrpA1-induced activation of just two dorsal projecting motor neurons, aCC and RP2, is sufficient to generate a "dorsal contraction" phenotype (RN2-Gal4 UAS-TrpA1; data not shown).

Anatomical characterization showed many lines that had dorsally-projecting motor neuron expression. Interestingly, there were lines that expressed in interneurons only and exhibited a similar phenotype (Figure 3C, 4). These interneurons are strong candidates for excitatory interneurons that directly or indirectly specifically stimulate dorsal-projecting motor neurons. We also found a line (R65D02) with muscle expression in dorsal acute and dorsal oblique muscle groups that gave a similar phenotype (data not shown).

#### Ventral contraction

We found 8 lines in this category; the R92C05 and R79E03 lines expressing only in sparse interneuronal patterns are shown in Figure 2I. Similar to the dorsal contraction phenotype, yet opposite in conformation, the ventral contraction phenotype was first discovered when we activated Gal4 patterns that expressed in ventrally projecting motor neurons (Nkx6, Hb9 and lim3B Gal4 lines; data not shown). When viewed laterally, the head and tail regions are ventrally contracted towards each other (Figure 2I’). Similar to the dorsal contraction postural phenotypes, we saw a spectrum of severity, with some continually stuck with tonically contracted ventral muscles, while others would go through bouts of ventral contraction, then can made attempts to crawl.

Anatomical characterization showed lines that had ventrally-projecting motor neuron expression. Interestingly, there were lines that expressed in interneurons only and exhibited a similar phenotype (Figure 3F, 4). These interneurons are strong candidates for excitatory interneurons that directly or indirectly specifically stimulate ventral-projecting motor neurons. We also found two lines (R40D04, R33E02) with muscle expression in ventral acute, ventral oblique and ventral longitudinal muscle groups that gave similar phenotypes (data not shown).

## Discussion

We identified a number of distinct behavior phenotypes elicited by activation of sparse subsets of neurons in the larval brain and VNC (Figure 5), but this is by no means an exhaustive exploration of TrpA1-induced larval phenotypes. As noted previously, roughly half of the statistically slow genotypes did not show any of the ‘overt’ phenotypic categories described in this paper. To fully characterize the remaining lines by phenotype would require advanced annotation of crawl dynamics and quantification of additional parameters. For example, upon high magnification observation of the slow hits, many simply appeared slow. Careful analysis by measuring wave duration and frequency may reveal additional phenotypes. Indeed, using refined analysis we investigated a slow line (R1 1F02) and discovered it was due to a defect in maintaining left-right symmetric muscle contraction amplitude during forward locomotion [19].

**Figure 5.**
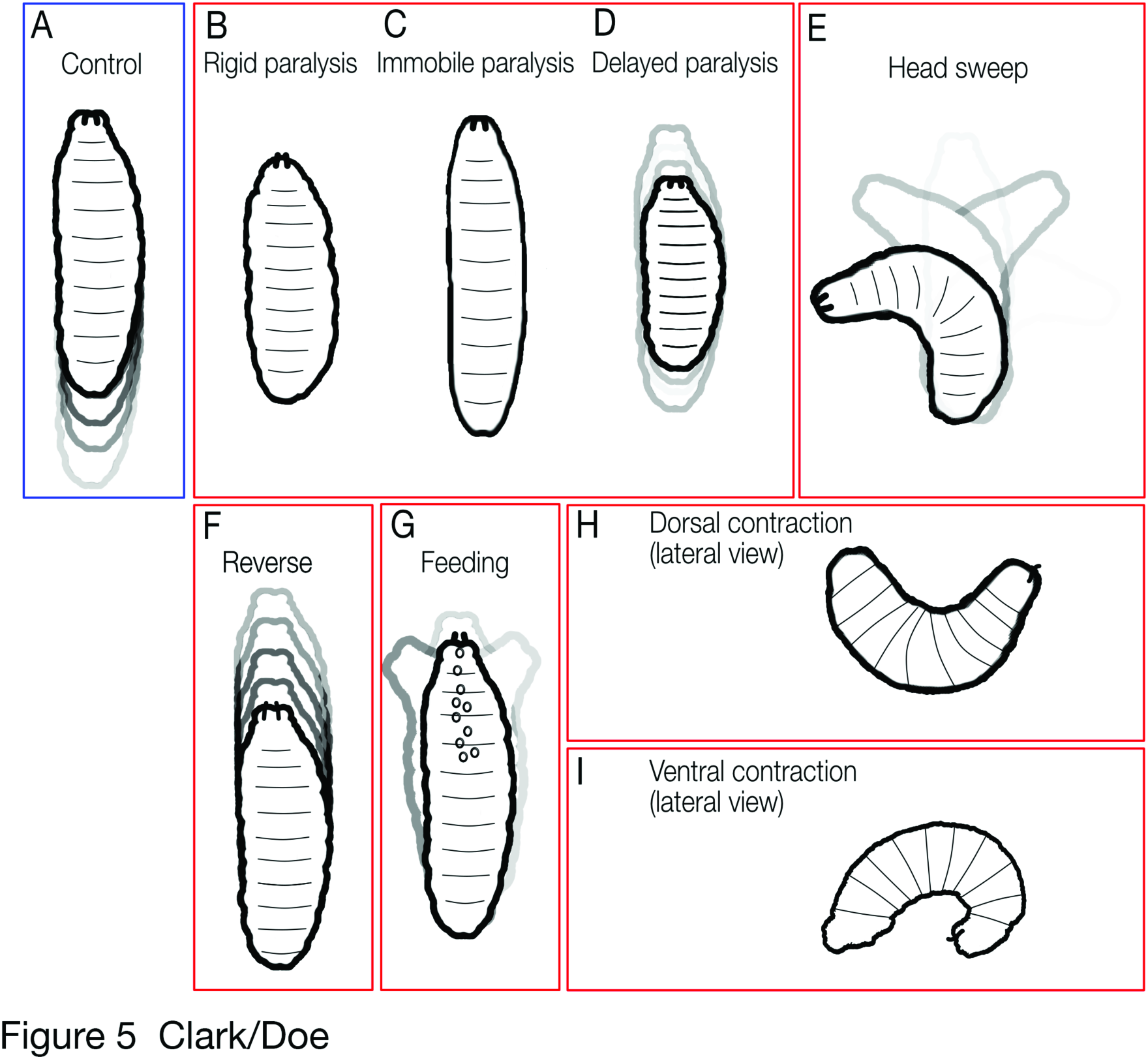
Summary of phenotypic groups. **(A)** Control larvae have free range of motion, crawling for bouts of forward or reverse (left, blue box). TrpA1-induced phenotypes bound in red (from left to right): **(B) Rigid paralysis:** complete loss of mobility with all segments of the larval body wall muscles fully contracted. **(C) Immobile:** complete loss of mobility with body wall segments often lacking tone, appearing smoothened and the larvae becoming languid and lengthened. **(D) Delayed paralysis:** gradual slowing of crawl speed over time until finally becoming immobile with tonic contraction of body wall muscles. **(E) Head cast:** head sweeps back and forth; can occur with thoracic/abdominal paralysis or with normal thoracic/abdominal peristaltic movements. **(F) Reverse:** only backward peristaltic movements. **(G) Feeding:** constant digging around with mouth hooks and attempts to ingest sub trate. Frequentrhythmic ingestion of gaseous bubbles can be observed. **(H) Dorsal contraction:** head and tail is raised off substrate. **(I) Ventral contraction:** head and tail are curled ventrally toward each other.

Recently developed larval tracking methods for multiplexed computational analysis would greatly assist the further definition of TrpA1-induced larval phenotypes. Examples of novel tracking methods include FIM, MaggotTracker, Multiple Worm Tracker and idTracker [34,58–60]. For example, MaggotTracker can characterize aberrations in run distance, duration, strides and many other quantitative parameters. This type of quantitative analysis can detect abnormal crawl patterns not readily identifiable by human eyes [34].

Many of the phenotypes we illustrated contained anatomical expression patterns with only interneurons, suggesting that those behavioral phenotypes were generated in the CNS. However, there were a large majority of lines that also expressed in tissues such as muscles, motor neurons, sensory neurons or glia. Many of these "off target" neurons can be discounted; for example, it is highly unlikely that motor neuron activation induces the head cast, reverse, or feeding phenotypes. Of course, motor neuron expression can lead to complex phenotypes, such as a combination of feeding + paralysis phenotypes (R21C06) or reverse + dorsal contraction phenotype (R53F07).

Some phenotypic categories contained single Gal4 lines, whereas some categories had multiple Gal4 lines that generated a particular behavior. The latter could be due to multiple lines expressed in a common neuron or pool of neurons -- or due to several different neurons being able to produce the same phenotype (e.g. premotor and motor neurons). Further characterization of the expression patterns of lines with similar phenotypes will be necessary to resolve this question.

In the future it will be important to define the neurons within each Gal4 line expression pattern that generates a specific motor pattern. *Drosophila* genetic techniques have made it possible to restrict expression of Gal4 patterns to successfully identify individual neurons that generate a behavior. For example, stochastic flipping [61,62], the FLP/FRT system [63,64], and the split-Gal4 system [65–67] all allow subdivision of a Gal4 pattern. An intersectional technique has used the FLP/FRT system to successfully dissect the functional elements of *the fru* circuit [63,68], and we recently used the split Gal4 system to identify a subset of functionally relevant interneurons governing muscle contraction amplitude during forward locomotion [19]. Application of these methods should allow identification of the neuron(s) responsible for each of the eight locomotor phenotypes described in this paper.

